# Pan-cell-type prediction of splicing patterns from sequence and splicing factor expression

**DOI:** 10.64898/2026.02.18.706660

**Authors:** Kalin Vetsigian, Jenessa Lancaster, Ioan Ieremie, Caleb M. Radens, Paul Smyth, Stephen Young

## Abstract

Alternative splicing is a core determinant of cell-type-specific gene expression in humans, and its dysregulation contributes to many diseases including neurodegeneration, autoimmunity, and cancer. However, current deep learning models for predicting RNA expression from sequence are limited in how they handle cellular context dependence. These models typically achieve cell-type specificity by training separate models or heads for each tissue or cell-type, which assumes discrete, predefined cell types. This design not only prevents learning from pathological or experimentally perturbed transcriptomes, but also prevents generalization to new cellular contexts. Here we introduce PanExonNet, a deep learning framework that integrates cis- and trans-regulation by conditioning sequence-based splicing predictions on an inferred “splicing state” derived from the expression of RNA-binding proteins (RBPs) and spliceosome components. PanExonNet is trained on diploid, individual-specific gene sequences containing indels to predict splicing profiles derived from short-read RNA-seq, and it outputs multiple tracks at single-nucleotide resolution as well as donor-acceptor junction usage. Compared to multi-headed baselines such as Borzoi and Pangolin, PanExonNet exhibits substantially higher context specificity—as quantified by a ΔPSI correlation metric designed to isolate context-specific variation—and, crucially, generalizes to unseen cell types. Adding perturbational knockdown followed by RNA sequencing (KD-RNA-seq) data to the training set further improves generalization. This performance is enabled in part by our introduction of contextualizable convolutions, a modular layer that may broadly benefit genomic sequence modeling. This framework provides a scalable foundation for future DNA-to-RNA models that could improve variant effect prediction, the design of oligonucleotide therapeutics, and biomarker discovery across diverse cellular contexts.

## Introduction

In eukaryotes, individual genes produce a diverse repertoire of mRNA isoforms in a regulated, cell-type-specific manner. A significant fraction of the isoform diversity is translated [1], resulting in protein variants with different functions, interactions and localization [2, 3, 4]. Other mRNA isoforms are byproducts of transcriptional noise [5] or players in a sophisticated layer of post-transcriptional control [6]. Strikingly, humans are among the species with highest isoform diversity [7], particularly in neural tissues [8, 9, 10], and dysregulation of the processes involved is causal or symptomatic of many diseases [11], including neurodegeneration [12], autoimmunity [13], and cancer [14, 15]. Correspondingly, there has been great interest in understanding healthy expression [16], detecting abnormalities [17, 18], and, ultimately, designing clinical interventions [19, 20].

Deep learning models, powered by vast genomic and transcriptomic datasets, have emerged as the primary tool for this task. While these models excel at learning complex cis-regulatory patterns from DNA sequence, they struggle to account for the dynamic, trans-regulatory environment that dictates cell-type specificity. The state-of-the-art approach—training separate models or heads for a finite set of tissues—imposes a rigid, categorical structure on what is biologically a continuous manifold of cellular states (Fig. 1a) [21, 22, 23]. This design choice is not only oversimplifying; it actively prevents learning from the most informative biological contexts, including idiosyncratic disease states, tumors, cell lines, and experimental perturbations that do not fit neatly into predefined categories. By treating cellular context as a discrete label rather than a continuous input, these models sacrifice the ability to generalize and are unable to leverage the full spectrum of available genomic and transcriptomic data.

**Figure 1:**
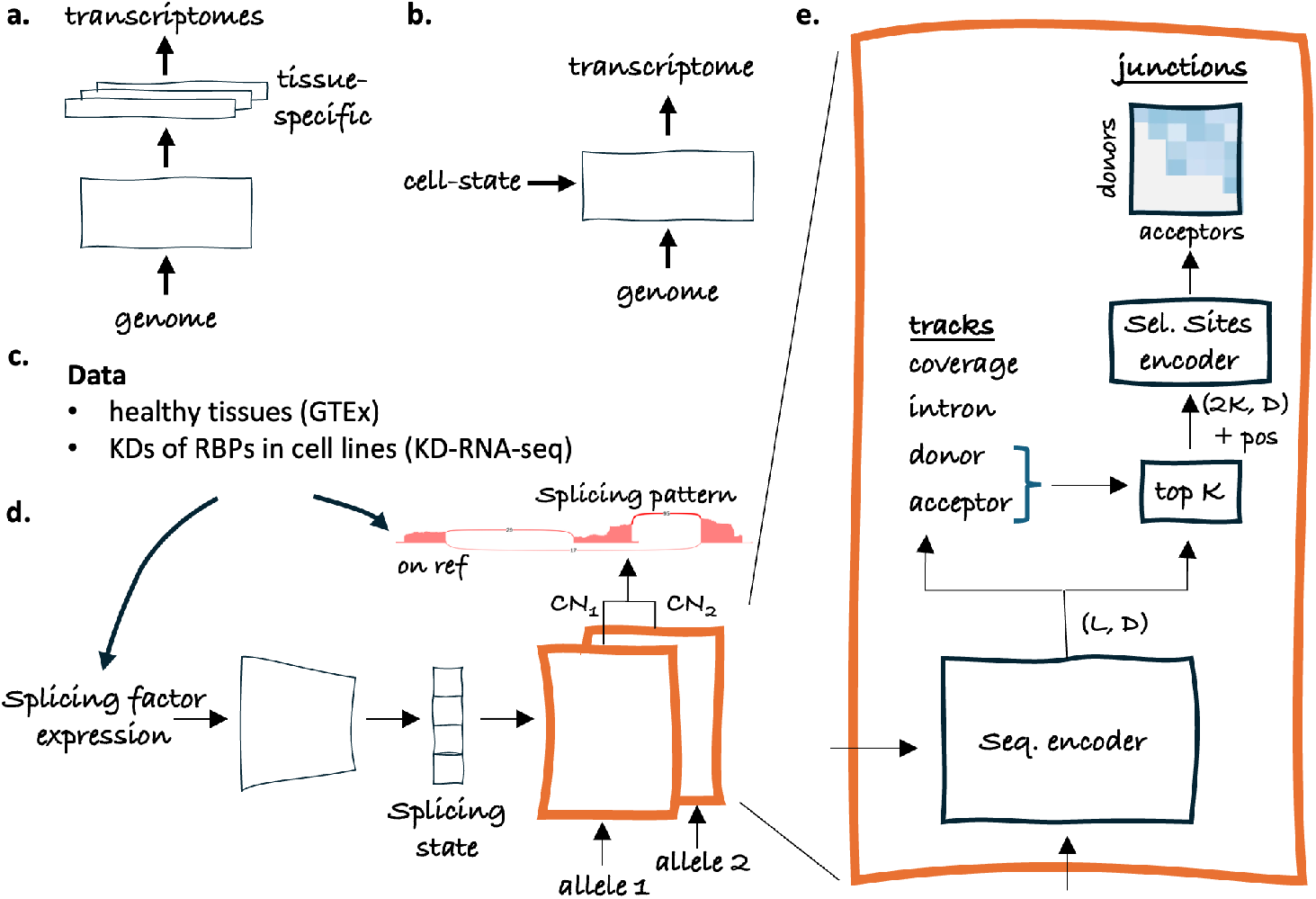
PanExonNet achieves cell-type specificity of splicing predictions from diploid or copy number-altered gene sequences via modulation from a cell state derived from splicing factor expression. **(a)** Most deep learning models that predict aspects of gene expression from DNA sequence achieve tissue-specificity by training a different head for each tissue. **(b)** An alternative is to use a cell-state embedding that captures trans aspects of gene regulation to modulate the genome to transcriptome map. **(c)** PanExonNet is trained on short read RNA-seq data from healthy tissues (GTEx v8) and knockdowns of RBPs in cancer cell lines (KD-RNA-seq). **(d)** PanExonNet infers a splicing state based on the expression of splicing factors and uses it to modulate the prediction of splicing profiles (sashimi plots) from gene sequences. Two copies of a genomic region containing variants and indels (along with their copy numbers in cell lines) are used as sequence input. Predictions are then combined by projecting them on the reference genome to match the standard RNA-seq processing practice. **(e)** A DNA sequence is fed into a contextualizable sequence encoder and followed by a head predicting four tracks at single nt resolution. The sequence embeddings of the top *K* donor and acceptor sites and their positions are then used to predict donor-acceptor junction usage.

Here, we advance an alternative approach in which the genome-to-transcriptome map is modulated by a relatively low-dimensional cell state, specific to each RNA-seq sample (Fig. 1b). While it is possible to treat this state as a latent variable that is learned for each sample, here we opt to learn it from the gene expression of a small number of selected genes—resulting in models that combine DNA sequence and gene expression inputs [24]. This allows training on any genome-transcriptome pair—independent of biological context— and learning representations of cell state and how they are controlled by the selected genes.

In this work, we restrict ourselves to predicting aspects of relative isoform distributions, as we are mostly interested in modeling RNA-processing post transcription initiation, and splicing in particular (Fig. 1d). By focusing on relative rather than absolute predictions about isoforms, we aimed to relieve the model, as much as possible, of the need to learn about the regulation of transcription initiation, which is likely much more complex and non-local than splicing, polyadenylation, and the other post-transcription processes.

The spliceosome is one of the largest and most complex molecular machines, and the expression levels of spliceosome components [25] and splicing factors are known to regulate or dysregulate global splicing patterns [26, 27, 28]. We therefore selected spliceosome components and splicing factors (mostly RNA binding proteins) as modulating genes and refer to the modulating cell state as a ‘splicing state’. Importantly, we only provide the model with the overall expression (transcripts per million, TPM) for those genes, while aiming to predict details about relative isoform expression for all genes. While conditioning on splicing factor expression has recently shown promise in predicting individual splicing features [29, 30], here we apply it to holistic DNA-to-RNA modeling.

The models presented here are trained on individual-level genomes (Fig. 1d, Methods). The sequence inputs are diploid gene sequences containing variants and indels. Predictions from the two gene copies are projected to the reference genome coordinates and combined, ensuring consistency with the standard practice of aligning RNA-seq reads to the reference genome. For cancer cell lines, we weigh the two gene copies depending on local ploidy, under the assumption that most heterozygous alleles were present before chromosome regions were replicated or lost. This handling of genomic variation goes above and beyond previous practices, Table 1. Most of the previous models use only reference genome sequences as inputs [31, 32, 21, 22], while those trained on individual-level data only utilized constitutional (germline) genomes [23].

**Table 1:** Comparison of models capable of multipurpose, tissue-specific DNA-to-RNA predictions.

|  | PanExonNet | Borzoi | Pangolin | AlphaGenome | BigRNA |
| --- | --- | --- | --- | --- | --- |
| Train on individual-level genomes | ✓ | x | x | x | ✓ |
| Handle copy number variation | ✓ | x | x | x | x |
| Predict donor-acceptor junctions | ✓ | x | x | ✓ | x |
| Predict overall expression of genes | x | ✓ | x | ✓ | ✓ |
| No cell-type annotations required | ✓ | x | x | x | x |
| Can learn from any genome-transcriptome pair | ✓ | x | x | x | x |
| Can generalize to unseen cell-types with gene-expression data | ✓ | x | x | x | x |
| Resolution of predictions | 1nt | 32nt | 1nt | 1nt | 128nt |

The models predict the relative isoform expression information contained in poly-A enriched short-read RNA-seq data. We extract coverage and split-read information from the short-read RNA-seq to effectively predict sashimi plots [33] (Fig. 1d,e). Specifically, we predict 4 tracks at single nucleotide (nt) resolution: coverage, donor usage, acceptor usage, and intron. The coverage track is derived from aligned RNA-seq reads, whereas the other three are different summarizations of information derived from split reads. Furthermore, we explicitly predict donor-acceptor junctions—not just splice sites (Fig. 1e). From the existing models, only the recently published AlphaGenome [21] has done that. Our approach to constructing the training objective differs in that we treat the sashimi plot as a coherent object, encompassing all tracks and junctions, which we try to predict up to an overall scale.

We refer to the resulting framework as PanExonNet. We compared different architectures within this framework and juxtaposed them with Borzoi (multi-headed) and Pangolin (multi-headed + tissue fine-tuned)^1^, as well as matching multi-headed models we trained. To demonstrate the framework’s ability to handle data from diverse contexts, we jointly trained on GTEx [16] and perturbation data in cancer cell lines from KD-RNA-seq [34]. We focused our evaluation on cassette exon inclusion, a downstream task not directly optimized during training that captures one of the primary mechanisms by which isoform distributions vary between cell types [35, 36].

## Results

### PanExonNet models with and without junction prediction and two modulation strategies were compared with each other and with multi-headed models

*Pan-track* models predict only tracks and have a sequence encoder based on novel contextualizable ConvNeXT layers we developed. *Pan-junct* models predict junctions in addition to tracks. In *pan-concat* models, the splicing state is concatenated to the sequence embedding after a ConvNeXT-based sequence encoder, and a channel-wise MLP is used to yield tracks. Multi-headed models have a ConvNeXT-based sequence encoder followed by tissue-specific heads. All models were trained on GTEx, but pan-junct models were also trained on GTEx + KD-RNA-seq. In each case, we trained several models and evaluated the ensembles (Methods).

### We constructed a cassette exon dataset to benchmark tissue specificity and generalizability

(Fig. 2a, Methods). Cassette exon inclusion is one of the primary ways in which isoform distributions vary between cell-types, which makes it particularly useful as a measure of tissue-specificity. Each exon is characterized by PSI (percent spliced in), which is equal to the fraction of mRNA molecules for a gene that include that exon. We compared several methods for extracting PSI information based on predicted coverage, splice-site strength, intron track, and junctions (Methods). For Borzoi, we only used coverage information since it is the only relevant track type predicted, and for Pangolin—splice-site strengths. We emphasize that none of the models are directly optimized for this task.

**Figure 2:**
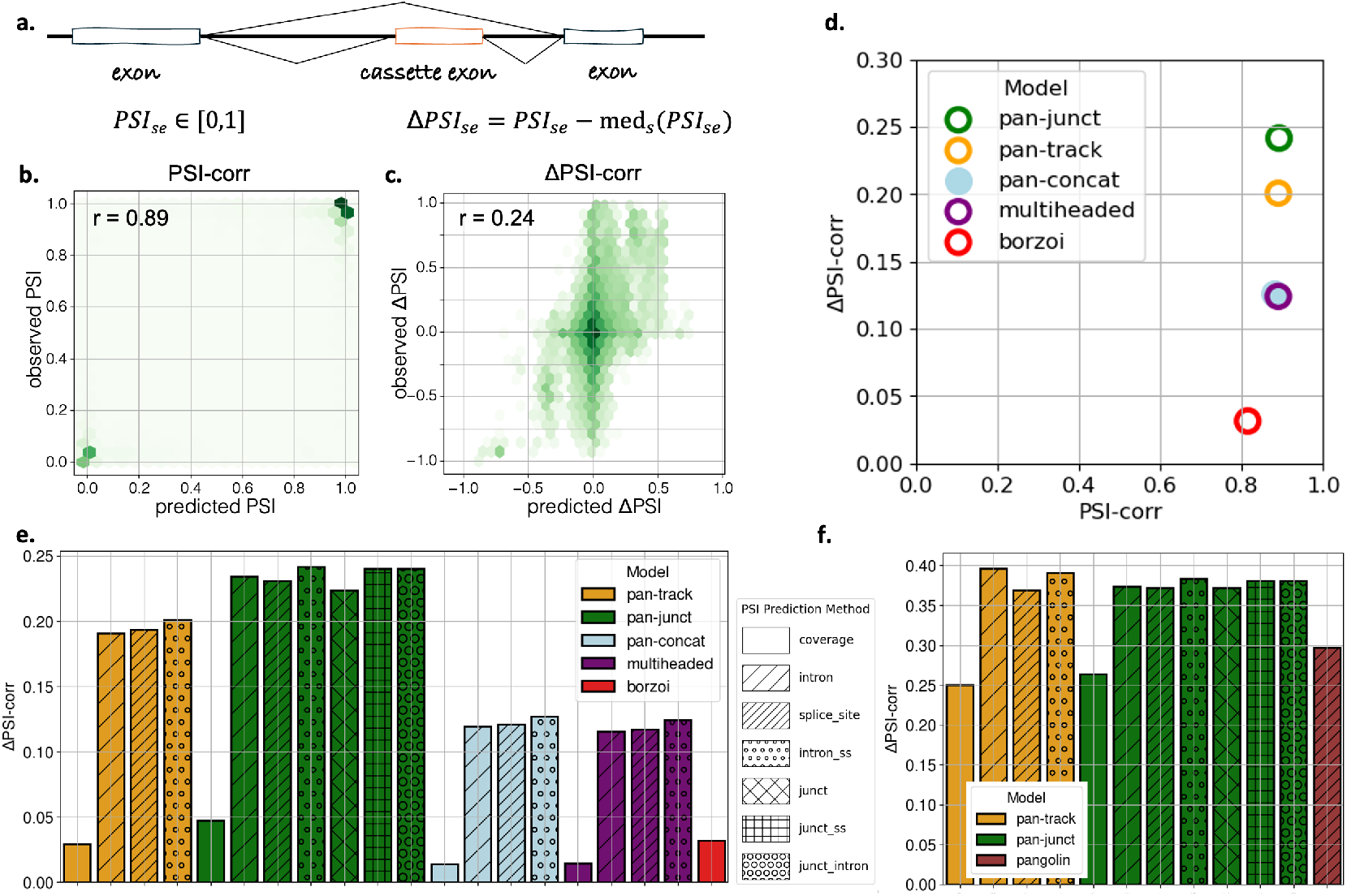
PanExonNet exhibits better tissue-specificity than models with tissue-specific heads or fine-tuning on a cassette exon inclusion benchmark. **(a)** The inclusion of a cassette exon *e* in a sample *s* is quantified by *PSI*_*se*_ ∈ [0, 1] (percent spliced in). Δ*PSI*_*se*_ is then defined as the deviation from the median *PSI*_*se*_ across training samples for that exon. **(b)** Scatterplot of predicted vs observed *PSI*_*se*_ for the pan-junct model. The *PSI*-*corr* metric is defined as the corresponding Pearson correlation. **(c)** Scatterplot of predicted vs observed Δ*PSI*_*se*_ for the pan-junct model. The Δ*PSI*-*corr* metric is defined as the corresponding Pearson correlation. **(d)** Comparison of five ensembles on these metrics: pan-junct, pan-track, pan-concat, multi-headed, Borzoi. While all models exhibit similarly high *PSI*-*corr*, they have much lower and differentiated Δ*PSI*-*corr* performances, indicating that Δ*PSI*-*corr* is a more useful metric of tissue-specificity. **(e)** Different models (colors) and different ways of deriving PSI from model outputs (hatching) are compared on Δ*PSI*-*corr*. **(f)** PanExonNet models with and without junction head are compared with Pangolin on the 4 tissues supported by Pangolin and the intersection of their held-out genes.

### We introduce ΔPSI correlation as a challenging and sensitive metric of context specificity

We first compared models in terms of direct correlation between predicted and observed PSI across contexts (Fig. 2b,d, Methods). We found that all models exhibit high correlation, indicating that all models can distinguish between typically included vs excluded cassette exons. To more sensitively measure context specificity, we examined the ability of each model to predict deviation from the median inclusion for each exon across samples in training tissues. Even a model that is good at predicting splicing features can score close to zero if the predictions are not cell-type-specific.

### PanExonNet models outperform Borzoi and Pangolin on tissue-specificity of exon inclusion

We performed a conservative comparison by evaluating on our held-out genes, whereas the Borzoi ensemble was trained on all genes. Even under this setup, the PanExonNet models achieve an order of magnitude improvement in tissue-specific exon inclusion prediction (Fig. 2d,e), despite using substantially lighter convolution-based sequence encoders compared to Borzoi’s transformer architecture. For the Pangolin comparison, we evaluated on the intersection of held-out genes for both models, restricting the analysis to the four tissues that Pangolin supports. These tissues have abundant training data across multiple species, a favorable scenario for tissue-specific models where no extra benefit from cross-tissue learning is expected. Nevertheless, the PanExonNet models exhibit superior performance (Fig. 2f).

### PanExonNet models are better than matched multi-headed models on tissue-specificity

Many of the major models achieve tissue-specificity by training tissue-specific heads [32, 23, 21, 22]. To better understand the advantages of the PanExonNet framework, we trained a multi-headed ensemble that mirrors pan-track in terms of architecture, outputs, and training data. We found that the pan-track ensemble has significantly higher tissue-specificity than the multi-headed ensemble on train tissues (Fig. 2d,e).

### Tracks informed by split-reads are superior for predicting tissue-specific exon inclusion than the coverage track

This trend is consistent across the different PanExonNet and multi-headed models (Fig. 2e,f). This can partially account for the lower performance of Borzoi, which only predicts coverage, and the higher performance of Pangolin, which predicts splice-site usage derived from split-reads. We also note the high performance of the novel intron track we introduced. In most cases, the top prediction method for PSI utilizes the intron track by itself or in combination with splice sites or junctions.

### A pan architecture based on contextualizable convolution is superior to concatenating the splicing state after the sequence encoder

Within the PanExonNet framework, there are different ways by which a splicing state can modulate the sequence encoder. We compared two approaches. One is based on novel contextualizable convolution layers, and the other concatenates the splicing state to the sequence embedding at each position before the predictive head. We found that contextualizable convolution (Fig. S2) leads to much higher performance on ΔPSI (0.2 vs 0.13, Fig. 2d,e) and several other metrics (not shown). The relative performance of different tracks for PSI prediction remains the same. Interestingly, pan-concat has similar performance to multi-headed, and both share the principle of a universal sequence embedding that is customized to different cell-types as late as possible. This suggests that differential processing from the ground up is a more powerful approach to impacting tissue-specificity. In addition, we found that concatenating or adding splicing state embeddings to the input does not lead to any tissue-specificity. While the space of potential PanExonNet architectures is large, contextualizable convolution appears to be a promising approach to contextualization of sequence encoders.

### Training to predict donor-acceptor junctions increases performance

Pan-junct models have higher performance than pan-track models (Fig. 2d,e). Strikingly, the addition of a junction head not only increased performance based on the predicted junctions but also boosted the ability of all tracks to predict exon inclusion. Thus, there is synergy rather than interference between the junction and track objectives.

### The junction head learns to predict non-trivial junctions beyond simply connecting neighboring splice-sites

The increased performance on cassette inclusion already suggests that the junction head learns non-trivial information. However, we can also explicitly ask whether it can predict non-trivial junctions given a knowledge of the splice-sites. Using the observed donor and acceptor sites from split reads in each sample and gene, we can separate junctions into classes of increasingly non-trivial order and, additionally, separate them by intron length (Methods). In every category that we examined, the ability to predict presence and absence of junctions significantly exceeded an expectation based on the mean junction frequency for that category (Fig. 3 and Fig. S4).

**Figure 3:**
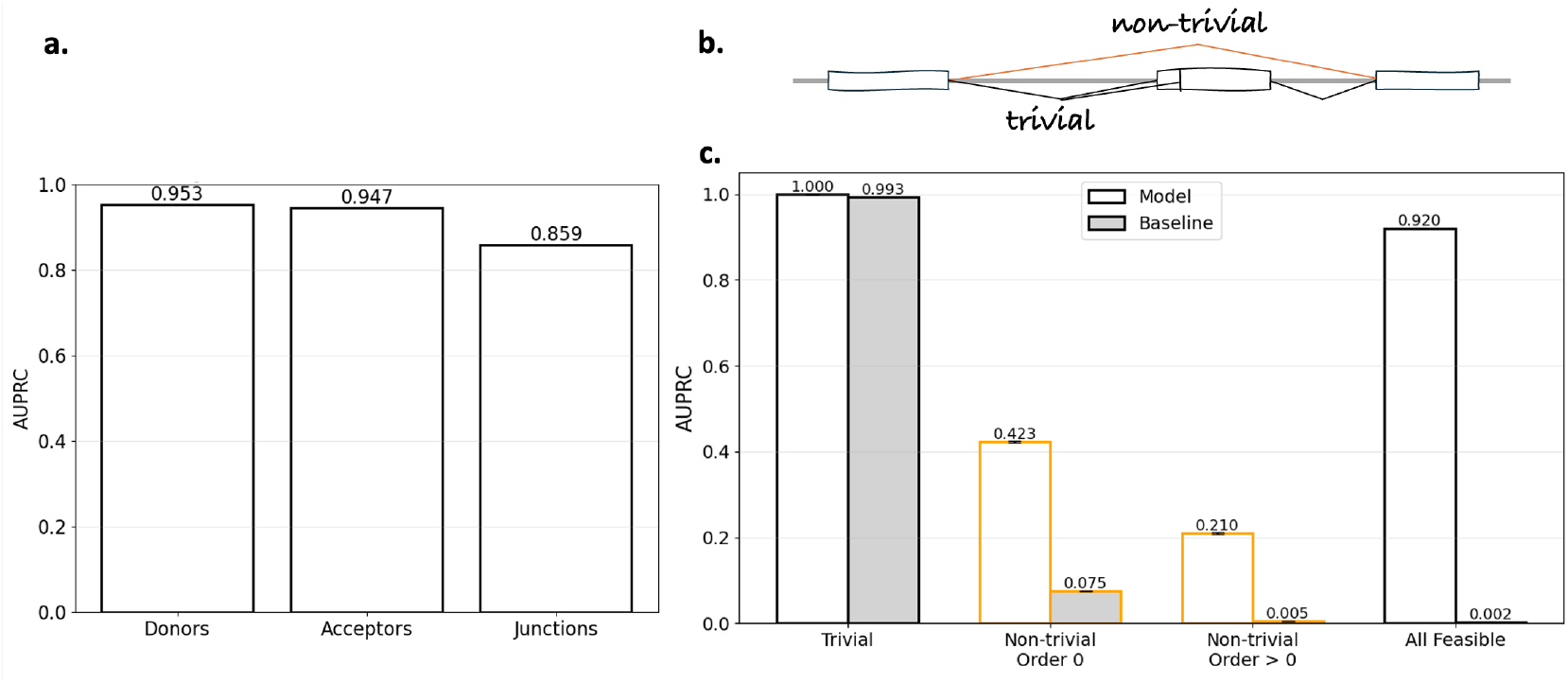
The junction head can predict non-trivial junctions. **(a)** AUPRC for prediction of donor sites, acceptor sites, and junctions in held-out genes. **(b)** Junctions can be classified into trivial ones—for which either the donor is connected to the nearest downstream acceptor, or an acceptor is connected to the nearest upstream donor—or non-trivial ones of various orders—by recursively removing the trivial ones (Methods). **(c)** AUPRC of the junction head on held-out genes, stratified by junction class, compared with a baseline that knows the donor and acceptor positions and predicts junctions at observed frequency for each class (Methods; see Fig. S4 for breakdown by intron length).

### PanExonNet models generalize to unseen cell-types

We trained pan-junct models by holding out two cell-types in GTEx with very distinct splicing factor profiles (Fig. 4a). We computed ΔPSI correlation on the held-out tissues and genes by subtracting the median PSI for each exon across train tissues. Note that negative ΔPSI correlation can occur if the model overfits to train tissue splicing factor expression and unseen combinations detune the sequence encoder, producing predictions worse than simply using the median values. Instead, we saw ΔPSI correlations that were of similar magnitude as those seen when looking at tissue specificity, indicating that the model exhibits ability to generalize to unseen cell types based on the expression level of splicing factors in those cell types.

**Figure 4:**
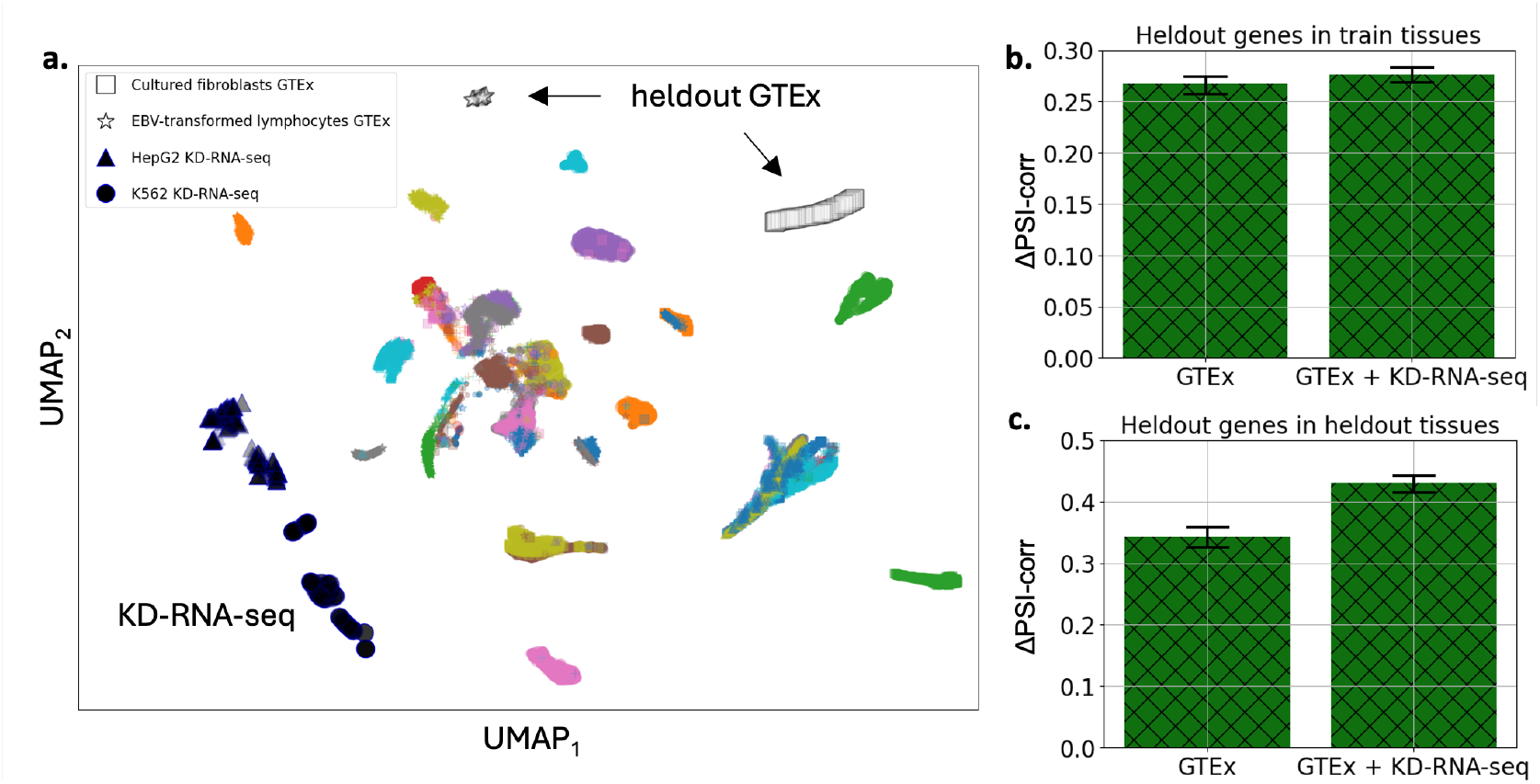
PanExonNet cassette exon predictions generalize to unseen cell types and are improved by training on intervention data from KD-RNA-seq. **(a)** UMAP embedding of splicing factor expression for samples in GTEx and KD-RNA-seq. Different colors and symbols are used for different tissues and cell-types. **(b)** Comparing Δ*PSI*-*corr* performance on held-out genes and train GTEx tissues of models trained on GTEx only or GTEx + KD-RNA-seq. The two models are compared on held-out genes and held-out GTEx cell-types. In both (b) and (c), Δ*PSI*_*se*_ was computed by subtracting the median over samples in the train set for each exon.

### Jointly training on GTEx and KD-RNA-seq improves generalization to unseen cell-types

One of the motivations for the PanExonNet framework was to develop an ability to jointly train on any available data, including cancer cell line data and perturbation data. The KD-RNA-seq dataset measures the effects of knockdown of RNA binding proteins on the transcriptomes [34]. Therefore, we asked whether adding this dataset to training data would improve the ability of the model to generalize to the unseen cell types. We saw that adding KD-RNA-seq slightly increases performance on train GTEx tissues, which we did not necessarily expect (Fig. 4b). But, importantly, it increased generalization to the held-out cell types (Fig. 4c). Thus, training on perturbation experiments in cancer cell lines increases performance on unrelated cell types, indicating the generalization power of the PanExonNet models.

### The models often fail to predict deviations from the median behavior but have good positive predictive value

Examining the ΔPSI correlations, we found that there are many false negatives for which the model predicted median behavior whereas the observed data points exhibited large deviations. However, wherever the model predicted large deviations, they were likely to be true. Correspondingly, if we ignore small ΔPSI predictions as unreliable, the metrics sharply increase, indicating that the model has few false positives (Fig. 5).

**Figure 5:**
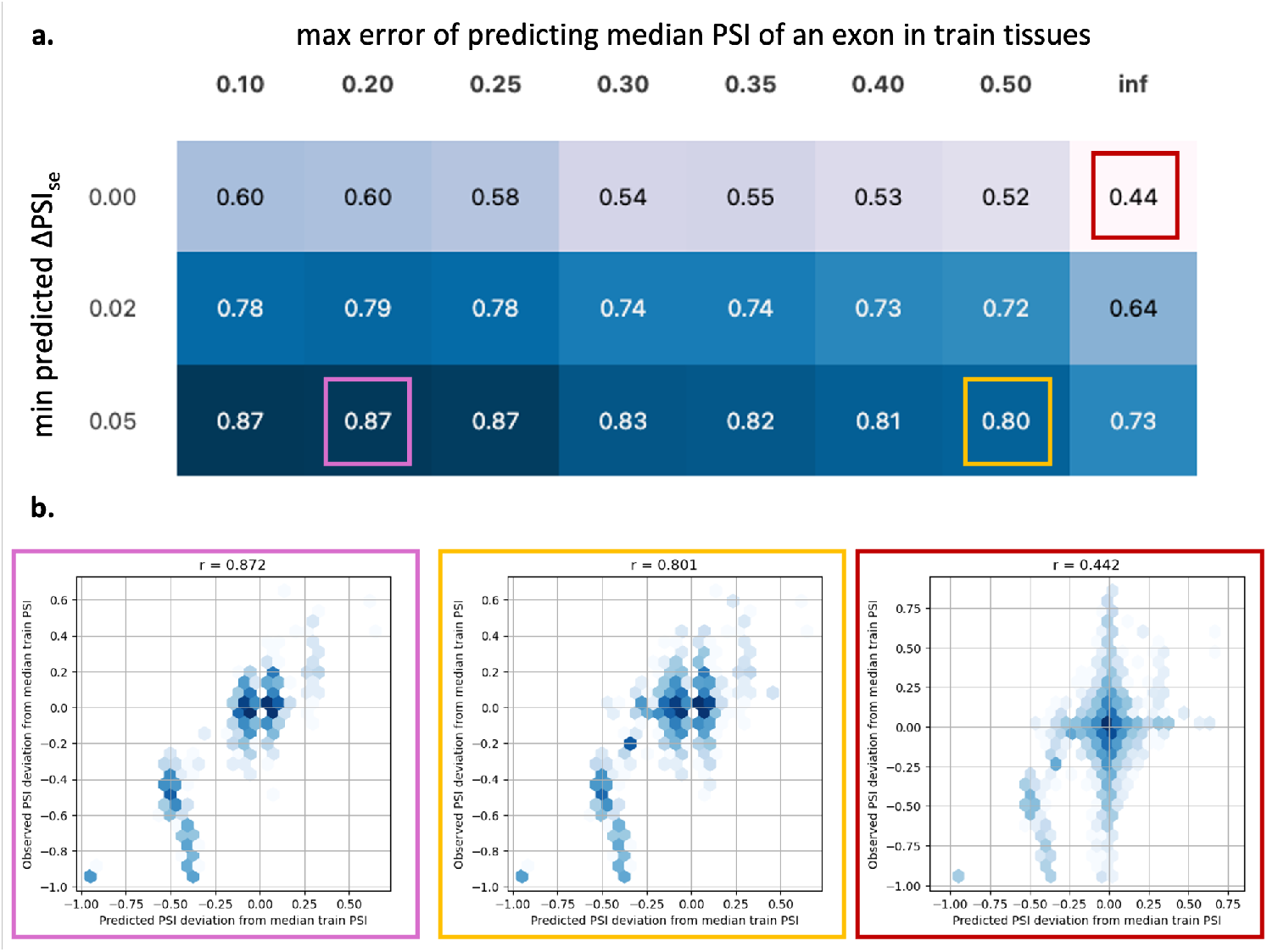
Large predicted deviations from the median *PSI* have a strong predictive value. Detecting and removing unreliable predictions can improve model usefulness. Shown here is the effect on Δ*PSI*-*corr* of removing instances with small predicted Δ*PSI*_*se*_ and/or exons for which the median predicted *PSI* over train samples is wrong. The results are for the GTEx + KD-RNA-seq model from Fig. 4, evaluated on the held-out cell types and genes. **(a)** Table cells display Δ*PSI*-*corr* values. Rows correspond to different thresholds that the absolute value of predicted Δ*PSI*_*se*_ needs to exceed to keep a data point, with the first row corresponding to no filtering. Columns correspond to different thresholds for removing exons based on the prediction error of their median *PSI* over train tissues, with the rightmost column corresponding to no filtering. **(b)** Scatter plots of predicted vs observed Δ*PSI*_*se*_ for some of the cells in (a). Frame colors match those of highlighted table cells.

Additionally, the models fail to accurately predict even the median behavior for a subset of exons. For each model, such exons can be identified based on their poor performance on the training tissues. We verified that excluding these exons from evaluation further improves the overall generalization performance in held-out tissues (Fig. 5a). Thus, by allowing the model to abstain in certain cases—either when predicting small deviations or when an exon exhibits poor training performance—we can substantially improve the reliability of the remaining predictions. The relatively high positive predictive value of filtered predictions enables downstream applications.

## Discussion

We developed a modeling framework to predict relative isoform expression from DNA sequence in a cell-type-specific manner. Cell-type specificity is achieved by providing the expression levels of a panel of splicing factors as input and inferring a splicing state that modulates the DNA-to-RNA mapping. This pan-cell-type framework can learn from any pair of human genome and transcriptome data without requiring additional annotations. This is distinct from most previous approaches for predicting molecular phenotypes from DNA sequence, which achieve tissue-specificity by training tissue-specific heads. A benefit of the pan-cell-type approach is that it can seamlessly integrate data from diverse healthy or pathological cell types and perturbation experiments without relying on predefined tissue or cell-type categories. This makes it conceptually possible to converge to a useful causal model of context-specificity in the limit of a lot of intervention data or through active learning. The framework also allows generalization to unseen cell types, which to our knowledge is a first-in-class capability for general-purpose DNA-to-RNA models.

Our modeling effort is also distinct in that it fully leverages the information contained in both short-read RNA-seq data and individual-level genomes (Table 1). The models were trained to predict all aspects of relative isoform expression that can be inferred from short-read RNA-seq, including the locations and usage of donor-acceptor splice junctions. Rather than relying on a reference genome, the models learn directly from individual-level genomic data, explicitly accounting for indels, diploidy, and even aneuploidy, thereby enabling training on and application to diverse cell lines and samples. To our knowledge, this is the first DNA-to-RNA model that trains on individual level data, predicts junction usage, and combines training on healthy tissues (GTEx) and perturbation experiments in cell lines.

Our work highlights the importance of benchmarks of context specificity, given that many machine learning models exhibit impressive performance at predicting complex molecular phenotypes from sequence but struggle to predict how changes of context or perturbations affect these phenotypes. It is often much easier to predict typical behavior than deviations from it. To benchmark tissue-specificity and generalizability in the context of splicing, we developed a cassette exon inclusion benchmark. The benchmark was constructed so that even models that are good at making non-trivial predictions of splicing patterns from sequence score close to zero if the predictions are not tissue-specific. Furthermore, models that are tissue-specific but overfitted to train tissues can score negative on generalizability. This benchmarking revealed large differences between different models and different ways of computing cassette exon inclusion from the predicted splicing patterns. It is yet unsaturated and can guide future modeling efforts.

We demonstrated significant improvements relative to Borzoi and Pangolin in terms of tissue-specificity of cassette exon inclusion. The improvements come from several factors. First, the PanExonNet framework is more performant than a matching multi-headed framework, particularly when the splicing state modulated all layers of the sequence encoder. Second, tracks derived from split-reads, i.e. the splice site usage and intron tracks, are more predictive than the coverage track, which is derived from aligning non-split mRNA reads. Third, training to explicitly predict junction usage increases performance when cassette inclusion is calculated from predicted junctions or tracks. Predicting junction usage also allows disambiguation of more complex splicing patterns such as mutually exclusive exons.

We also demonstrated generalizability to unseen cell types for which gene expression data exists, ensuring that the held-out cell types were sufficiently different. We further showed that adding training on perturbation data in which RNA binding proteins are knocked down in cancer cell lines (KD-RNA-seq) improves generalization performance— demonstrating the benefits of training on data from diverse cell-type contexts within the PanExonNet framework. Generalizability implies that perturbation experiments in cell lines and accessible tissues can be used to learn about splicing regulation and predict interventions in clinically relevant cellular contexts. Such generalizability can be used, for example, to improve disease diagnosis by predicting splicing variation in clinically nonaccessible tissues [37] or to predict neoantigens resulting from aberrant splicing in particular tumor types [14].

A major motivation for this work was to develop an approach optimized for modeling RNA-processing post transcription initiation based on short-read RNA-seq data. Past efforts have focused on predicting absolute expression levels, which has the advantage of being holistic. But we opted for predicting relative isoform expression, which factors out the complex and non-local regulation of transcription initiation under the approximation that in most cases there is no strong coupling between transcription initiation and splicing. Such an approximation has potential benefits, given the current quality, diversity, and quantity of transcriptomic data, compared with approaches that aim to predict the full transcriptome. A systematic comparison is left for future work.

Many genomic sequence models use convolutional layers in the initial layers of their sequence encoders, as they are known for efficiently learning binding motifs and other local features. Contextualizable convolution can be used to make them context specific in a straightforward manner. It is a computationally efficient way to confer specificity, which can, moreover, be implemented so that genome-wide inference for a specific cell-type has zero computation overhead relative to a regular convolutional model. Biologically, much of cell-type specificity comes from changes in the abundance or state of proteins with different binding preferences, and the resulting effective change in the relative importance of sequence motifs can be efficiently modeled as context dependent weighing of different filters. We based our implementation on ConvNeXt because it is a strong convolutional architecture that does not require batch normalization and is therefore compatible with small batch sizes, which allows the use of longer sequences. Thus, contextualizable ConvNeXt is a modular layer that can broadly benefit genomic sequence modeling tasks where context-specificity is important.

A biological limitation of our approach is that the splicing state, i.e. the collection of trans determinants of cell-type-specific splicing, is not fully determined by the expression of RNA-binding proteins and splicing factors. For example, mutations of splicing factors are ignored, which is particularly relevant for cancers [15, 19]. Additional regulatory layers, such as post-translational modifications (e.g., phosphorylation of splicing factors), are also not captured by expression levels alone. The assumption that the splicing state is fully determined by expression can be relaxed in the future by including a sample-specific latent component.

Another assumption is that cell-type specificity is encoded in trans, i.e. that it can be represented by a global and relatively low-dimensional state. Epigenetic modifications of DNA are, of course, cell-type-specific and can affect transcription in cis, which is not explicitly modeled. To the extent that relevant cell-type-specific epigenetic differences cannot be inferred from a global state, the modeling cannot capture them. This limitation is shared with the multi-headed approach in the typical case where the tissue-specific head has a relatively small number of parameters. However, our modeling makes the additional assumption that we can infer any relevant global determinants of epigenetic modifications from the particular panel of genes chosen for modulation. Our panel included putative splicing factors, but it will be interesting to experiment with inclusion of global regulators of cell-type-specific epigenetic modifications. Our framework is also compatible with the addition of epigenetic tracks to the input.

The splicing state inferred from the expression of splicing factors can be useful in its own right. It is a compact representation of the aspects of cell state relevant for global RNA processing. The splicing state can be used to identify biomarkers of disease states that are reflected in altered RNA processing. Furthermore, the context encoder can be used to predict and design interventions that shift the splicing state in a desired direction.

The contextualization by splicing factor expression can be used as an interface to single-cell data, which typically only measures gene expression. By inferring the splicing state for each cell, it becomes possible to predict splicing patterns in single cells based on their splicing factor expression. This can be useful for understanding cell-type-specific splicing patterns in complex tissues and developmental trajectories.

This work is a proof of concept and can benefit from scaling up. Most importantly, it can be extended to encompass additional datasets such as ENCODE [38] and CCLE [39], which is expected to greatly increase generalization performance. Cell lines have many more mutations than what is available within the germlines, which can also improve variant effect prediction. Taking advantage of long-read data can further increase performance. We deliberately kept a convolution-based sequence encoder to emphasize that the PanExonNet framework is an orthogonal direction of improvement to scaling. However, scaling the sequence encoder with attention layers could be beneficial, even though our selected-sites encoder and junction head already brings in some of the benefits of long-distance attention in an efficient manner. The number of genes used to infer the splicing state can also be increased. Here, only 277 splicing factors were used, but there are many more RNA-binding proteins [40] and global cell-type-specific regulators [41] that could be worth including. Finally, the ability of the network to integrate data from different contexts allows systematic improvements through lab-in-the-loop active learning via perturbation experiments on different cell-type backgrounds.

## Methods

### Models

There are two main PanExonNet model variants: *pan-track* models predict tracks only, whereas *pan-junct* models predict both tracks and junction usage. For both model types, the context encoder, sequence encoder, and track head are the same. The *pan-junct* models have two additional components: selected-sites encoder and junction head. Two additional model varieties were used in comparisons: pan-concat and multi-headed, which output only tracks.

#### Inputs

Inputs are provided as batches of *B* entries. During training, each entry consists of two allele sequences along with information aligning them to the reference sequence and an expression vector containing TPM values for a panel of *G* genes (subject to augmentation). For cell lines, a copy number is also provided for each allele to capture aneuploidy. The sequences are of length *L* + 2*F*, where *L* is the length of the core region over which predictions are made, and *F* is the length of flanking regions on either side. The alignment information for each allele sequence is provided as a vector of length *L* containing the indices within the reference gene sequence to which each position within the core region maps. Insertions are represented by repeating the same reference index for the length of the insertion rather than advancing it, whereas deletions are represented by skipping reference indices in the alignment vector. The alignment information is important because long sequences are likely to contain some indels and because the RNA-seq data used for training has been mapped to the reference genome.

Allele sequences are DNA letters encoded as one-hot vectors of size 4 (A, C, G, T) and (0, 0, 0, 0) for padding (P). The sequences are always provided in the direction of transcription – they are reverse complemented for genes on the negative strand. During inference, the model can be used with a single sequence input, e.g. for variant effect prediction.

#### Outputs

The model outputs a splicing profile consisting of 4 tracks and, for junction models, a junction usage matrix. The four tracks are: coverage, donor usage, acceptor usage, and intron. The coverage track is trained to be proportional to the read coverage in RNA-seq data along the gene, i.e. the number of reads overlapping a given position. The donor, acceptor, and intron tracks are learned from split-read data, providing different summaries. The donor/acceptor usage track aims to be proportional to the number of split reads with a donor/acceptor at a given position, whereas the intron track represents the sum of all split reads that skip a given position. Junction usage is represented as a vector of donor coordinates, a vector of acceptor coordinates, and a matrix of junction usage values for each donor-acceptor pair. Matrix elements corresponding to an acceptor coordinate less than the donor one (in direction of transcription) are ignored. Junction usage aims to be proportional to the number of split reads supporting each junction. Moreover, the correct relative scales of the different tracks and junction usage are learned during training, so that the model can output all components together as a consistent object determined up to an overall scale. The outputs taken together constitute a splicing profile similar to a sashimi plot (for a fraction of a gene).

#### Combining allele outputs

The output is obtained by projecting the allele-specific predictions (with the same gene expression context input) to reference coordinates using the alignment information and then averaging the two allele-specific outputs. For cell lines, the averaging is weighted by the copy number for each allele. Only predictions from the core region of length *L* are used for each allele. Junctions that start or end outside the core region are not returned. To resolve insertions when projecting to reference, the donor and acceptor usage tracks were summed over the insertion positions before projecting to reference (to preserve potential splice sites), whereas the track values over insertion positions were unused for coverage and intron tracks. We note that both insertions and splice-site positions are rare (on per nt basis), and therefore the resolution method of the exceedingly rare cases in which there is an insertion over a splice-site is not expected to impact training dynamics. Deletions relative to reference resulted in zero values along the deletion for all tracks. The returned donor or acceptor coordinate vectors for junction usage are a union of the selected donor or acceptor sites for each allele after projection to reference coordinates. Junction usage values are then averaged across alleles, with missing junctions (present in one allele but not the other) treated as zero. The length of the returned, merged tracks can differ slightly from *L* due to indels.

#### Loss function

All tracks and junction usage values are concatenated into a single vector per sample to form a splicing profile. The loss is based on a weighted cosine similarity between the predicted and target splicing profiles. To weight different components (tracks, junction matrix) of the splicing profile, each component’s values are multiplied by the square root of its assigned weight in both the predicted and target profile before computing cosine similarity. Any padded positions in the input sequence are ignored, and the contribution to the total loss is weighted by the fraction of unpadded positions. The loss contribution was not weighted by the number of reads supporting the target profile, to avoid giving excessive weight to highly expressed genes. The weights used were: coverage track: 1.0, donor usage track: 10.0, acceptor usage track: 10.0, intron track: 1.0, junction usage matrix: 100.0.

##### PanExonNet architecture

The architecture is overviewed in Fig. 1d,e. A context encoder is used to process the gene expression vector into a context embedding. Each allele sequence is processed separately through a shared sequence encoder that is modulated by the context embedding. A track head processes the sequence embeddings to output the four splicing tracks after removing the flanking regions.

For junction models, the top *K* scoring donor and acceptor sites within the core region (as determined from the donor and acceptor usage tracks) and their coordinates are processed by the selected-sites encoder (based on relative position attention) to produce splice site embeddings. Finally, the junction head processes the splice site embeddings to output a *K × K* matrix of junction usage values. *K* = 100 was used.

The two allele-specific outputs are then projected to reference coordinates and combined as described above.

#### Contextualizable ConvNeXt block

It is based on the ConvNeXt block architecture [42], which consists of a depthwise convolution followed by LayerNorm, a pointwise convolution that increases the number of channels by a factor of 4, a non-linear activation, and another pointwise convolution that returns the number of channels to the starting value. To make the block contextualizable, we adopted ideas from [43] and modulated its parameters based on the input context (Fig. S2). The context embedding is first passed through an MLP to derive a set of modulating parameters. These parameters are then split into groups and used to modulate the weights of the depthwise convolution, normalization, pointwise convolution, and the final channel weighing as shown. We used *h* = 1 heads in the depthwise convolution for all experiments in this work.

#### Sequence encoder

The sequence encoder architecture is shown in Fig. S1 and is a tower of contextualizable ConvNeXt blocks grouped into superblocks with skip connections. It takes as input a batch of sequence embeddings and a context embedding and outputs sequence embeddings. The depthwise convolutions were dilated, and the total receptive field was 10k nucleotides as in [31]. 128 channels were used.

#### Context encoder

The context encoder processes the gene expression vector to generate a splicing state embedding. We used a minimal context encoder: an MLP with one hidden layer of dimension 32 followed by a ReLU and final context embedding of size 16.

#### Track head

The track head is a pointwise fully connected layer that reduces the number of channels to 4, representing the four splicing tracks.

#### Selected-sites encoder

(Junction models only.) The embeddings for the top *K* scoring donor and acceptor sites within the core region (based on the donor and acceptor usage tracks) and their coordinates were processed by attention layers. The attention layers used relative positional information as described in [44, 45] to take into account the distance between splice sites. The positional embeddings were constructed from asymmetric relative distance basis functions using exponential, central mask, and gamma functions. These attention layers were incorporated into standard transformer blocks: layer normalization and multi-head attention with a skip connection, followed by a layer norm and a two layer MLP with another skip connection. We used 4 transformer blocks with 16 heads, key size of 64, value size of 128, and 128 channels.

#### Junction head

(Junction models only.) A multi-head cross-attention layer with relative positional embeddings is used between keys derived from donor embeddings and queries derived from acceptor embeddings to generate an attention matrix of shape (K, K, H), where H is the number of attention heads. This is followed by an MLP operating on the last dimension to output the junction usage matrix of shape (K, K). *H* = 16 and MLP with one hidden layer of size 4 was used. The attention layer parameters were the same as in the selected-sites encoder.

##### Pan-concat model

The sequence encoder consists of regular ConvNeXt blocks without context modulation. The splicing state embedding is passed through an MLP, concatenated to the sequence embeddings before or after the sequence encoder and projected back to regular number of channels. The track head was extended by adding two pointwise hidden layers of size 32 and 16, with LayerNorm and GeLU activations after each, before the final output layer leading to 4 tracks.

##### Multi-headed model

The context encoder was removed, and in the sequence encoder, contextualizable ConvNeXt blocks were replaced by regular ConvNeXt blocks. The track head was replaced by multiple tissue-specific track heads.

#### Ensembling

Ensembling improved performance for all models. It was done by averaging the predictions for tracks and junctions. Because the train loss was scale free, the typical magnitude of tracks and junction usage varied between checkpoints. To give each checkpoint an equal weight within an ensemble, we normalized the outputs from each checkpoint by dividing them by their scale estimated from inference over many genes and tissue contexts. Ensembles of 5 checkpoints were used for the models used in Fig. 2 and Fig. 3. Ensembles of 3 checkpoints were used for Fig. 4 and Fig. 5.

### Data

#### GTEx data

The GTEx data [16] used for the analyses described in this manuscript were obtained in Dec 2019. Individual genomes and alignment chains were extracted from the following phased vcf file provided by GTEx: `GTEx_Analysis_2017-06-05_v8_WholeGenomeSeq_838Indiv_Analysis_Freeze. SHAPEIT2 phased.vcf.gz` Number of reads covering each genome position and number of split reads supporting each junction were extracted for each sample from RNA-seq BAM files (`GTEx_Analysis_2017-06-05_v8`). hg38 was used as the reference genome for both DNA and RNA sequences.

#### KD-RNA-seq data

High-resolution aneuploid maps and mega haplotypes for HepG2 and K562 cell lines were obtained from [46] and [47]. They were used to distribute variants from ENCFF713BPG (HepG2) and ENCFF752OAX (K562) linked-read phased vcf files to long haplotypes and assign a copy number to each. 4.2% of K562 and 7.4% of HepG2 variants were randomly assigned due to lack of sufficient phasing information. We trained only on gene regions along which the copy number for both alleles was constant. RNA-seq data (BAM files) for the KD-RNA-seq experiments in HepG2 and K562 cell lines were obtained from [34]. hg19 was used as the reference genome for both DNA and RNA sequences.

#### Training data preparation

Gene sequences were chunked into overlapping segments of length *L* + 2*F*, with a stride of *L/*2, making sure that every gene region was within some core input region. Only gene sequences between beginning and end positions of the longest annotated transcript (from GENCODE26) were used, with padding symbols added before and after as needed to reach length *L* + 2*F* . Regions of gene overlap were not included in the loss to avoid ambiguities. For each sample, only gene regions with at least one junction supported by at least 25 split reads were retained to eliminate splicing profiles with poor coverage. *L* = 5000 and *F* = 5000 were used for training the non-junction models and *L* = 16000 and *F* = 5000 for the junction models.

#### TPM augmentation

During training, the cell type context data was augmented by replacing each gene’s TPM value, *T*_*g*_, with *T*_*g*_ *c/C*_*g*_, where *C*_*g*_ denotes the raw read count for that gene in the sample and *c* ∼ Pois(*C*_*g*_), thereby accounting for the sampling noise inherent in read count data and reducing overfitting to particular expression values. Genes with zero read counts (*C*_*g*_ = 0) were kept at zero TPM without augmentation.

#### Data splits

Only protein-coding genes were used for training. The same 20% of genes, selected at random, were held out during training across all models. All results presented are for held-out genes. For the results of Fig. 4, the held-out “tissues” were: Cultured fibroblasts and EBV-transformed lymphocytes.

#### Jointly training on GTEx and KD-RNA-seq data

To jointly train on GTEx and KD-RNA-seq data, we randomly sampled batches from GTEx and KD-RNA-seq with a ratio of 4:1.

#### Panel of splicing factors

We used the list of splicing factors assembled in [48]. We removed 3 genes that were not present in the GTEx expression data, resulting in a final list of 277 splicing factors.

### Cassette exon benchmarking

Binary cassette exon identification: aligned RNA-seq BAM files from GTEx v8 were quantified for alternative splicing with MAJIQ build and MAJIQ PSI (each tissue built separately, all settings default except MAJIQ PSI –min-experiments 1), and splicing events characterized per tissue with VOILA modulizer with default settings [49]. Binary cassettes were filtered from the cassette.tsv modulizer output by restricting to complex=FALSE modules. Only cassettes with both a source and target LSV were retained to remove possible spurious cassette exon events. For Fig. 2, we excluded GTEx tissues that we could not match from Borzoi coverage tracks. For comparisons with Pangolin, we used only the 4 tissues supported by the Pangolin model.

#### Methods for extracting PSI from model outputs

Individual DNA sequences, centered on the cassette exon, were extracted from each GTEx sample and used to generate splicing profile predictions across tissues. For each cassette exon, we extract the average coverage over the cassette exon (*C*_*CE*_) and its two flanking exons (*C*_1_, *C*_2_), the average intron track over the two flanking introns (*I*_1_, *I*_2_) and the cassette exon(*I*_*CE*_), the donor and acceptor splice site strengths of the cassette exon (*A*_*CE*_, *D*_*CE*_) and the flanking exons (*D*_1_, *A*_2_), and the three junction usages – non-skipping (*J*_1_, *J*_2_) and skipping (*J*_12_) as shown in Fig. S5. These features are then used to compute PSI using different methods:

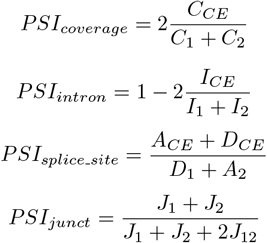

All values were clipped to the range [0, 1]. We also computed the average of junction and splice-site based PSI, junction and intron based PSI, and splice-site and intron based PSI. For Borzoi, we used *PSI*_*coverage*_ because coverage is the only relevant track type predicted. We used the 89 GTEx tracks and the mappings provided by the authors. For Pangolin, we used *PSI*_*splice_site*_ based on their splice-site strength tracks.

### Splice site and junction evaluation

The splice-site and junction AUPRC metrics in Fig. 3a are for random chunks from held-out genes from random GTEx samples. Splice-sites and junctions supported by at least 1 high-quality split read were counted as positives.

### Non-trivial junction benchmarking

We segregated junctions into increasingly non-trivial classes for benchmarking in Fig. 3c and Fig. S4. We start with the sets of observed donor and acceptor splice sites within a region. We define the class of feasible junctions as all pairs of donor and acceptor sites for which the acceptor is downstream of the donor. Within this class, we define trivial junctions as those for which the acceptor is the nearest acceptor downstream of the donor or the donor is the nearest donor upstream of the acceptor. We can now continue recursively by removing all trivial junctions from the set of feasible junctions and repeating the process. We define the trivial junctions obtained in the second round as non-trivial junctions of order 0. The rest of the junctions are non-trivial junctions of order *>* 0.

Within each class, we compute AUPRC and use the observed junction frequency as a baseline. In addition, within each class, we remove from consideration any potential junctions for which a splice site was not among the top *K* predicted donors or acceptors by the sequence encoder. This was done for ease of implementation and is the reason why the *All Feasible* metric in panel (c) is higher than the *Junction* metric in panel (a). In other words, panel (c) shows the performance of the selected-sites encoder and junction head only without penalizing splice sites missed by the sequence encoder.

Fig. S4 further breaks down the non-trivial junctions of order 0 by the distance between the donor and acceptor site. In every category that we examined, the predictions beat the observed junction frequency baseline.

## Acknowledgements

We thank Kim Branson for suggesting and supporting work on splicing modeling and Ron Schwessinger for valuable feedback on the manuscript.

---

**Supplementary Figure S1:**
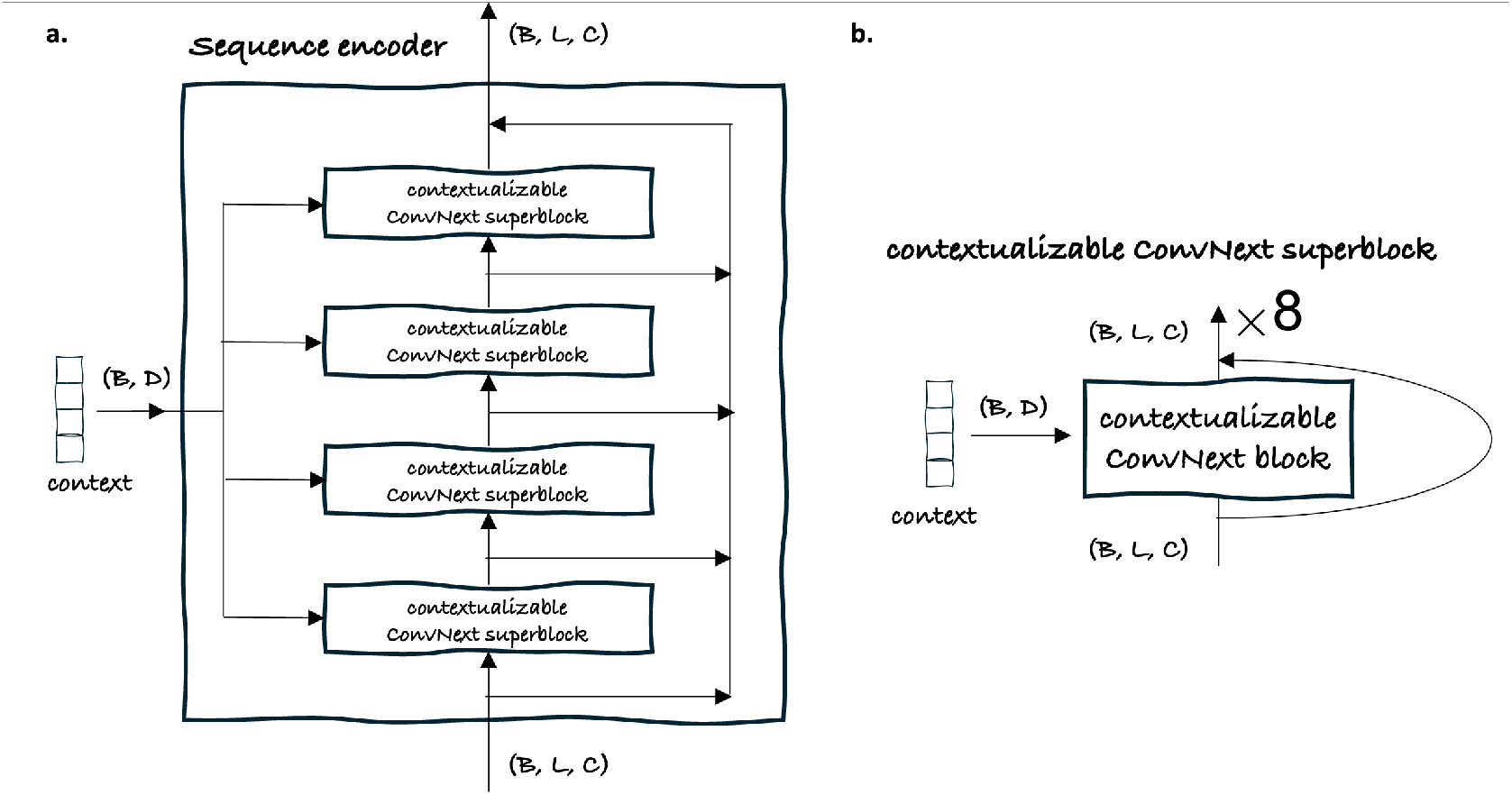
Sequence encoder architecture. Four contextualizable superblocks, each consisting of eight contextualizable ConvNeXt blocks, were used.

**Supplementary Figure S2:**
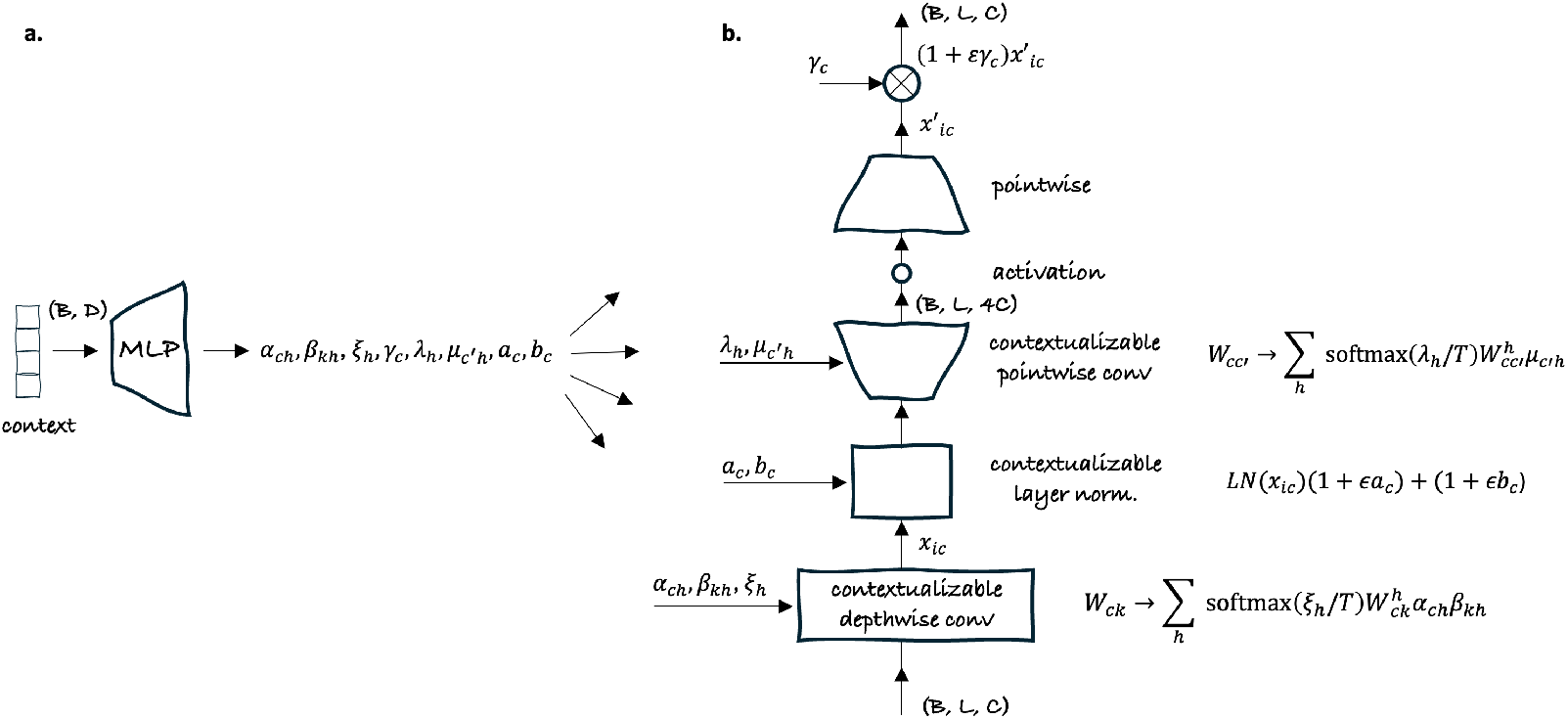
Contextualizable ConvNeXt block architecture. A batch of contexts of shape (*B, D*) modulates the processing of a batch of sequence embeddings of shape (*B, L, C*). For simplicity, the batch index is suppressed in all formulas. The small letter indices *c, h, k, c*^*′*^ and *i* enumerate, correspondingly: *C* channels, *H* heads, convolution kernel size *K*, 4*C* expanded channels, and *L* sequence positions. **(a)** The context is passed through an MLP and projected into several modulating parameters. This provides a layer-specific context processing. **(b)** The primitives of a ConvNext block are contextualized as explained by the formulas on the right.

**Supplementary Figure S3:**
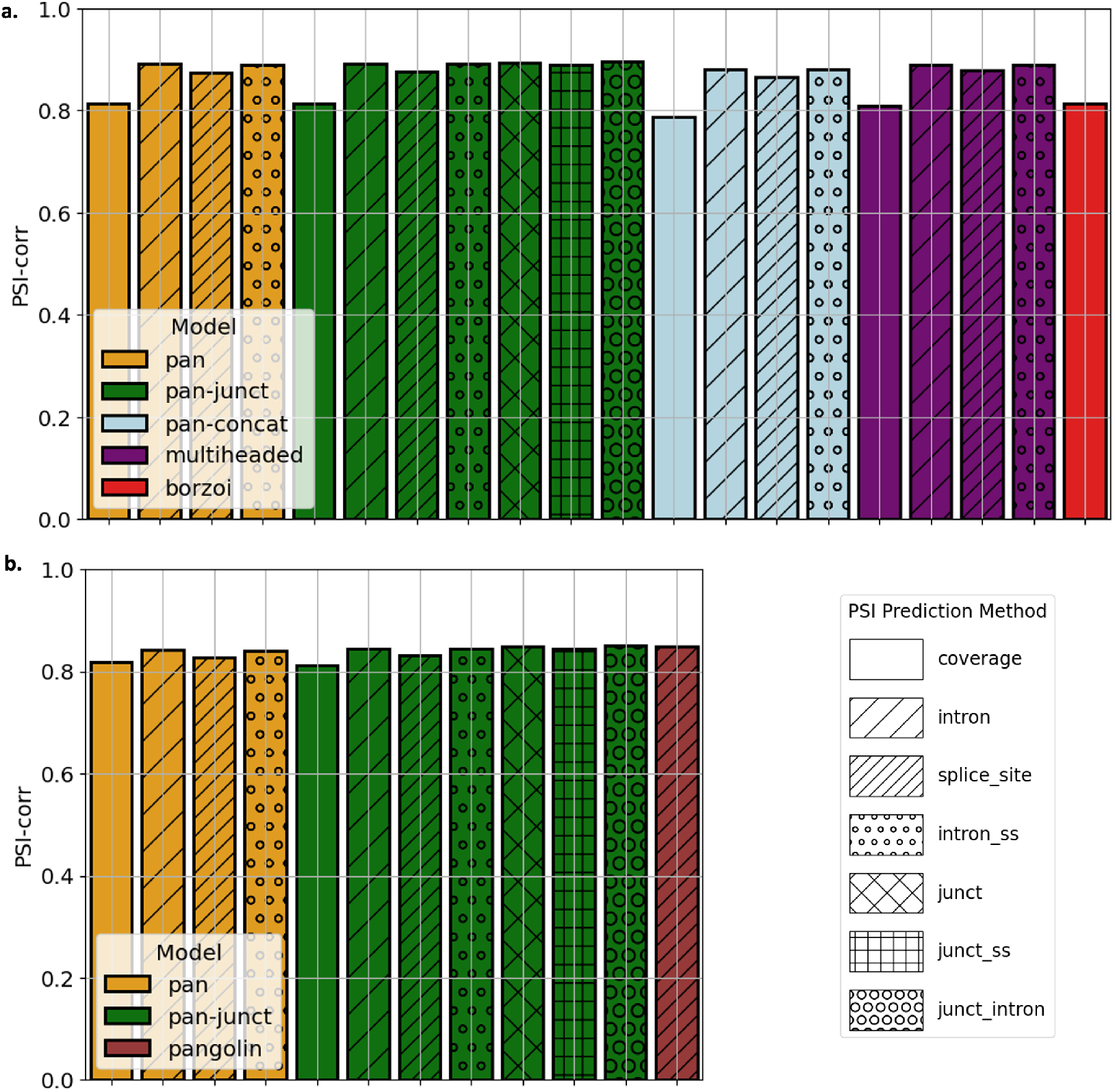
PSI-corr for different models and methods for predicting PSI. **(a)** PSI-corr for the models presented in Fig. 2e. **(b)** PSI-corr for the models presented in Fig. 2f.

**Supplementary Figure S4:**
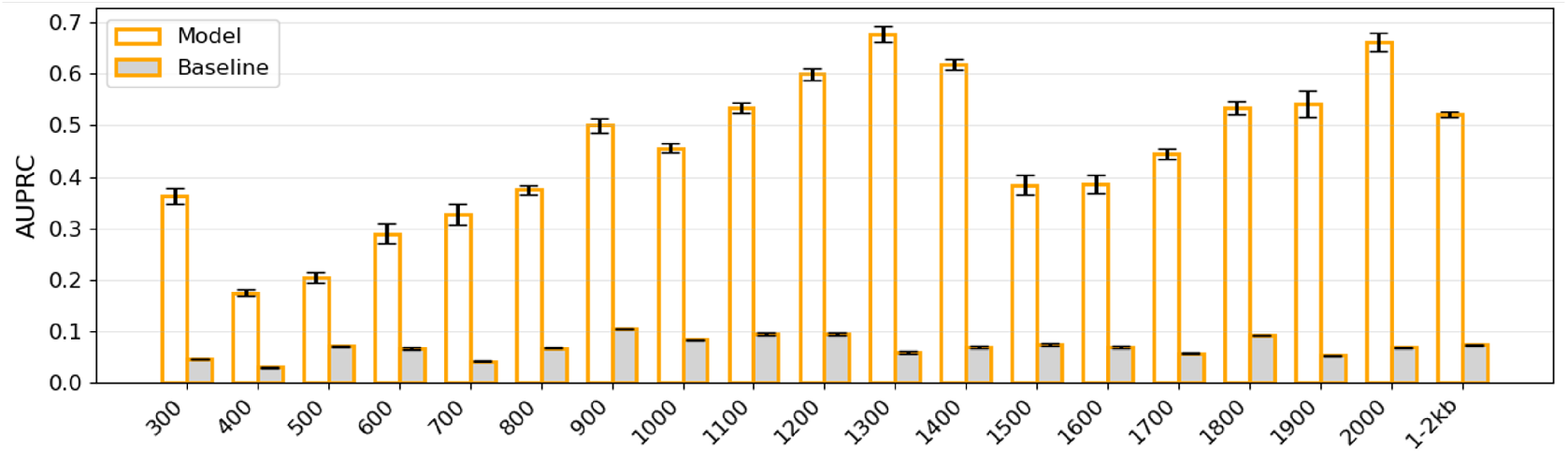
Prediction performance for non-trivial junctions by distance between donor and acceptor sites. AUPRC for the non-trivial junctions of order 0 from Fig. 3c segregated by distance between donor and acceptor sites. The junction head’s performance is compared with a baseline based on the average observed frequency (Methods). All but the last bin are of size 100 nt with the number beneath indicating the end of the bin, i.e. the first bin is 200–300 nt.

**Supplementary Figure S5:**
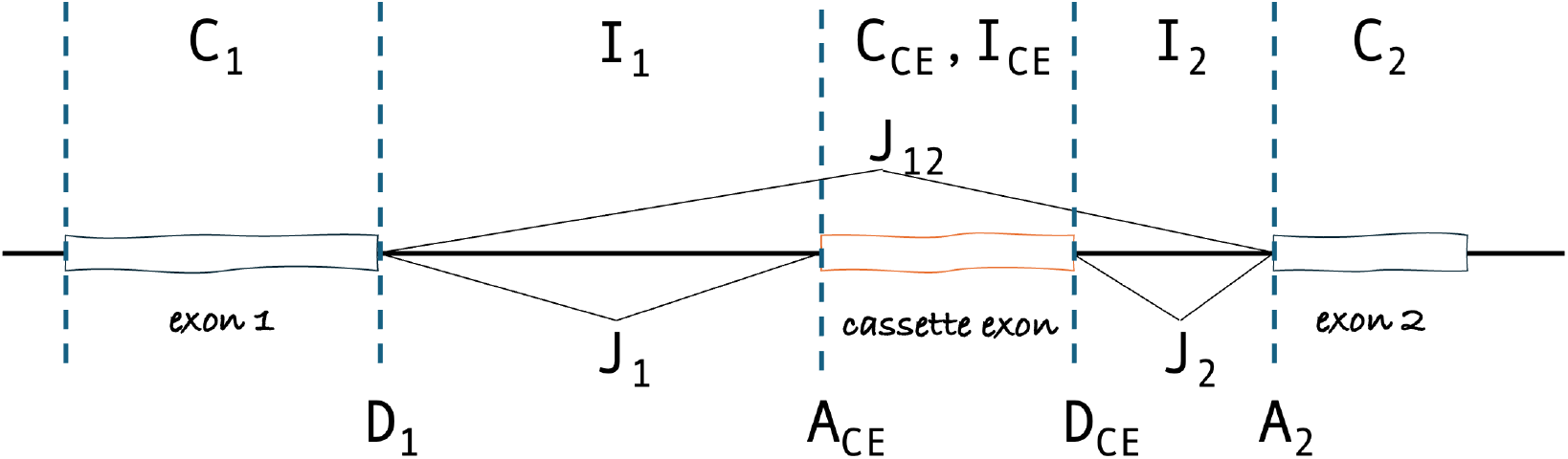
Extracting features from splicing profiles for cassette exon inclusion estimation. For each cassette exon, we extract the average coverage over the cassette exon (*C*_*CE*_) and its two flanking exons (*C*_1_, *C*_2_); the average intron track over the two flanking introns (*I*_1_, *I*_2_) and the cassette exon (*I*_*CE*_); the donor and acceptor splice site strengths of the cassette exon (*A*_*CE*_, *D*_*CE*_) and the flanking exons (*D*_1_, *A*_2_); the three junction usages—non-skipping (*J*_1_, *J*_2_) and skipping (*J*_12_). These features are then combined in different ways to compute PSI (Methods).

## Footnotes

1 AlphaGenome and BigRNA did not exist or have publicly available weights at the time of this research.

